# Brain enlargement with rostral bias in larvae from a spontaneously occurring female variant line of *Xenopus*; role of aberrant embryonic Wnt/β-catenin signaling

**DOI:** 10.1101/2023.10.23.563695

**Authors:** Ikuko Hongo, Chihiro Yamaguchi, Harumasa Okamoto

## Abstract

Increased brain size and its rostral bias are hallmarks of vertebrate evolution, but the underlying developmental and genetic basis remains poorly understood. To provide clues to understanding vertebrate brain evolution, we investigated the developmental mechanisms of brain enlargement observed in the offspring of a previously unrecognized, spontaneously occurring female variant line of *Xenopus* that appears to reflect a genetic variation. Brain enlargement in larvae from this line showed a pronounced rostral bias that could be traced back to the neural plate, the primordium of the brain. At the gastrula stage, the Spemann organizer, which is known to induce the neural plate from the adjacent dorsal ectoderm and give it the initial rostrocaudal patterning, was expanded from dorsal to ventral in a large proportion of the offspring of variant females. Consistently, *siamois* expression, which is required for Spemann organizer formation, was expanded laterally from dorsal to ventral at the blastula stage in variant offspring. This implies that the active region of the Wnt/β-catenin signaling pathway was similarly expanded in advance on the dorsal side, as *siamois* is a target gene of this pathway. Notably, the earliest detectable change in variant offspring was in fertilized eggs, in which maternal *wnt11b* mRNA, a candidate dorsalizing factor that activates Wnt/β-catenin signaling, had a wider distribution in the vegetal cortical cytoplasm. Since lateral spreading of *wnt11b* mRNA, and possibly that of other potential maternal dorsalizing factors in these eggs, is expected to facilitate lateral expansion of the active region of the Wnt/β-catenin pathway during subsequent embryonic stages, we concluded that aberrant Wnt/β-catenin signaling could cause rostral-biased brain enlargement via expansion of *siamois* expression and consequent expansion of the Spemann organizer in *Xenopus*. Our studies of spontaneously occurring variations in brain development in *Xenopus* would help uncover genetic mutations that drive analogous morphogenetic variations during vertebrate brain evolution.

## 1. Introduction

One of the most remarkable features of vertebrate evolution has been encephalization, characterized by an increase in the ratio of brain to body size. The degree of encephalization is quite substantial, with typical differences of 10-fold and 2-3-fold between reptiles and birds, and between birds and mammals (Martin, 1981). In some species, encephalization manifests itself primarily as neocorticalization, an increase in the ratio of the size of the cerebral cortex, the most rostral part of the brain tissue, to the total brain size. Extensive neocorticalization has played a prominent role in the evolution of primates, especially humans. The human cerebral cortex is approximately 3.5 times larger in weight (de Sousa and Wood, 2007) and three times larger in number of neurons, even when compared to our closest phylogenetic relatives, chimpanzees (Herculano-Houzel and Kaas, 2011). Recent investigations have increasingly focused on the genetic variation between humans and chimpanzees (Yousaf et al., 2021), with some interesting findings. These include increased proliferation of neurogenic cells attributed to human-specific *Arhgap11b* (Florio et al., 2015; Heide et al., 2020) and increased density of longer spines attributed to human-specific *Srgap2* (Charrier et al., 2012), both observed in the human neocortex. However, the striking differences between humans and chimpanzees in overall brain structure are not yet fully understood. Furthermore, our understanding of the mechanisms governing the evolution of the vertebrate brain as a whole remains incomplete. We may need a broader perspective and experimental approaches that better incorporate analyses of the cellular and molecular mechanisms involved in determining the overall structural arrangement of brain tissue in vertebrate species. Interestingly, in this context, adult populations of wild guppies (*Poecilia reticulata*) have been found to exhibit phenotypic variability in brain size relative to body size, with large-brained female lines generated by artificial selection showing higher cognitive performance than small-brained lines (Kotrschal et al., 2012). Anatomical analysis has shown that brain enlargement is rostrally biased and largely confined to the telencephalon, the most rostral part of the adult fish brain (Fong et al., 2021). However, the developmental and genetic mechanisms of this telencephalic enlargement have not been elucidated.

In the present study, we investigated the developmental mechanisms responsible for brain enlargement in larvae from a spontaneously occurring female variant line that we unexpectedly found in a farmed *Xenopus laevis* population. Variant females of this line produce offspring with varying degrees of enhanced dorsal and anterior structures (dorsoanteriorization phenotypes: Class I - III, defined in *2.1.*, Fig. 1A). Among these offspring, those with a relatively mild phenotype (Class I) showed evidence of brain enlargement with a rostral bias at the larval stage. Interestingly, the spectrum of dorsoanteriorization phenotypes observed in these variants closely resembled a range of phenotypes previously produced by artificial embryonic manipulation, including brain enlargement (Kao and Elinson 1988; Scharf et al., 1989). However, the unexpected finding that spontaneously variant females produced a significantly increased number, if not all, of their offspring with characteristically enlarged brains aroused our interest in investigating the underlying developmental mechanisms, with the prospect that the results might shed light on encephalization in vertebrate evolution. The rationale for this prospect stems from the notion that the mechanisms governing neural development, particularly in the early stages, are fundamentally conserved across extant vertebrates, including *Xenopus*, and are therefore likely to be shared with ancestral vertebrates. A prime example of this conservation is the induction of the neural plate and its initial rostrocaudal patterning, which is mediated by specific embryonic tissues commonly known as organizers (Houston, 2017; Stern, 2005; Wilson and Edlund, 2001). The neural plate then progressively subdivides to form the basal neural structures along the rostrocaudal axis common to vertebrates: the forebrain, midbrain, hindbrain, and spinal cord (Murakami, 2017). These basal structures give rise to further subdivisions with distinct cellular architecture and function characteristic of different vertebrate species. For instance, in mammals, especially humans, the cerebral cortex, which arises from the most rostral part of the forebrain, is prominently developed (neocorticalization). Mutations that led to the evolutionary divergence of the brain between vertebrate species could have occurred at any stage along this continuum of neural development, but in general, mutations at earlier stages of development may have had a greater impact on divergence.

**Fig. 1.**
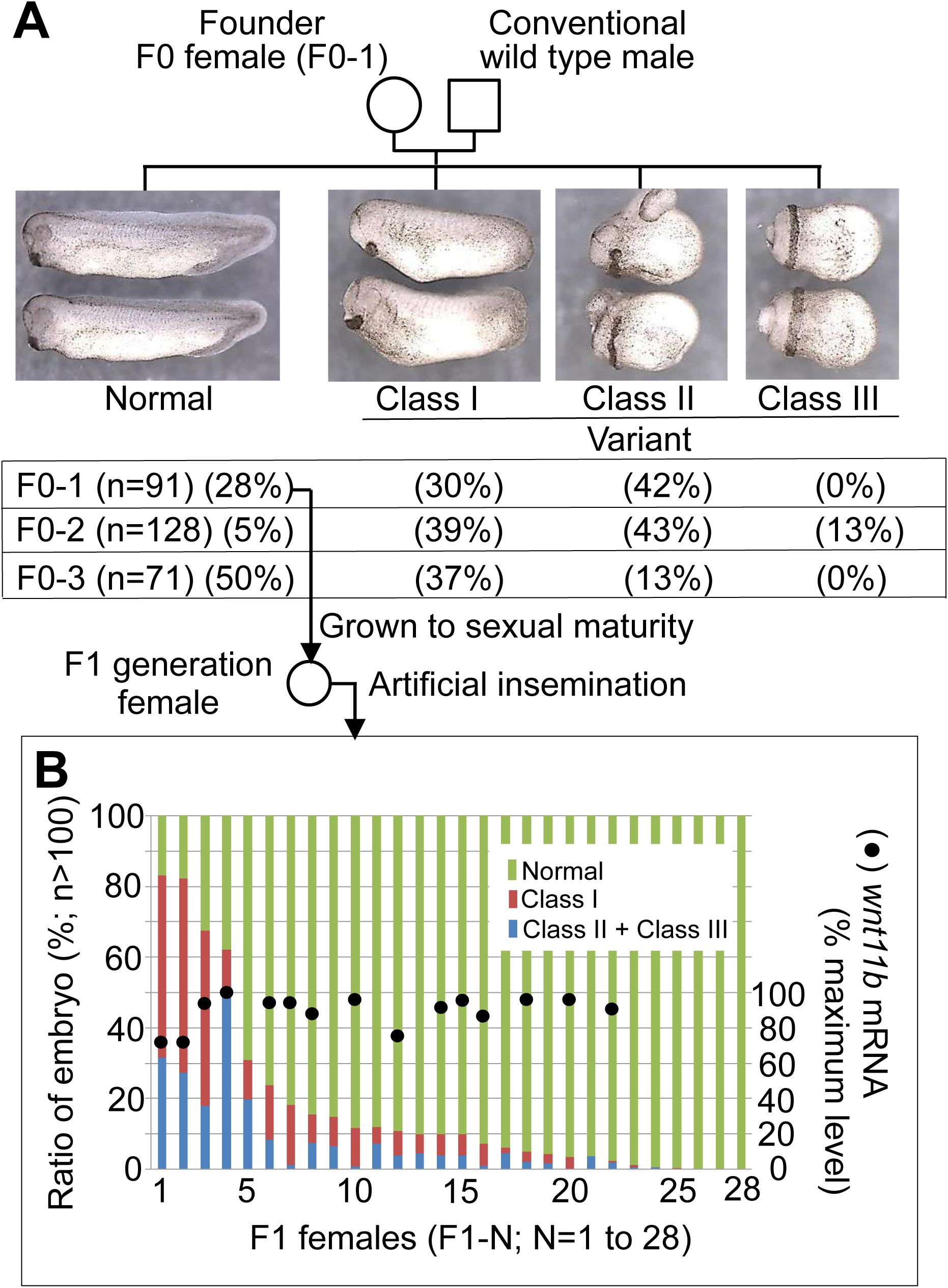
Genealogy of spontaneously variant females of *Xenopus laevis*. (A) Morphology of F0-1 larvae classified at stage 29/30. The table shows the proportions of each class in larvae from F0-1 to −3. n is the total number of larvae examined in a mating. (B) The proportions of each class in larvae from F1-1 to −28 (colored bars). The solid circles on the bars show the levels of *wnt11b* mRNA in unfertilized eggs of the respective F1 females, quantified from the data in Fig. 8A.

*Xenopus* proved advantageous for the purposes of the present study because, starting with the seminal Spemann-Mangold experiment (Spemann and Mangold, 1924), a wealth of data has been collected on the cellular and molecular mechanisms that govern early neural development in amphibians. In particular, a variety of marker gene probes have been successfully employed in recent years, some of which allow detailed analysis of organizer development and function, while others allow differentiation and tracing of individual brain subdivisions from the time of their formation. Notably, most of these *Xenopus* genes have homologs in the mouse genome; they appear to play similar developmental roles in these divergent species, as inferred from their expression patterns during early embryogenesis (Xenbase, MGI database).

In this study, using whole-mount *in situ* hybridization and serial sectioning, we showed that the brains of larvae with a relatively mild phenotype (Class I) were enlarged at the tailbud and tadpole stages. The degree of enlargement was more pronounced toward the rostral side, especially in the forebrain region. The rostral-biased enlargement was observed as early as the formation of the brain primordium, as indicated by the expression patterns of marker genes specific for the entire neural plate or its subdivisions. Remarkably, in a large proportion of gastrulae from variant females, the normally dorsally localized Spemann organizer was expanded ventrally to varying degrees, as evidenced by the expanded expression of *chordin* (Sasai et al., 1994) and *cerberus* (Bouwmeester et al., 1996). These genes encode proteins that are secreted from the Spemann organizer and initiate rostral neural induction. Another significant finding in the variant offspring populations was the dorsal-to-ventral (lateral) expansion of *siamois* expression at the blastula stage, again to varying degrees. Siamois is an essential transcription factor for the activation of *chordin* and *cerberus* in *Xenopus* (Engleka and Kessler, 2001; Kessler, 1997). Since *siamois* expression is induced by prior activation of the Wnt/β-catenin pathway (Brannon et al., 1997), the lateral expansion of *siamois* expression reflects a similar pattern of expansion of the active zones of Wnt/β-catenin signaling at earlier stages of development. Consistently, we also found that the distribution pattern of maternal *wnt11b* mRNA was expanded in the majority of fertilized eggs from variant females. In normal development, maternal *wnt11b* mRNA initially localizes to the vegetal cortical cytoplasm (VCC) of laid eggs and migrates dorsally soon after fertilization with other potential maternal dorsalizing factors. However, in many of the fertilized eggs of variant females, this maternal *wnt11b* mRNA was more diffusely distributed in the VCC compared to the control, with varying degrees of distribution. Since Wnt11b has been implicated in *siamois* expression via Wnt/β-catenin signaling (Brannon et al., 1997; Tao et al., 2005), it is most likely that the lateral spreading of *wnt11b* mRNA distribution contributes to the ventral expansion of *siamois* expression by laterally expanding the active domain of the Wnt/β-catenin pathway, resulting in the ventral expansion of the Spemann organizer. The relatively mild phenotype of rostral-biased brain enlargement observed in some of the variant larvae (Class I) is thought to be a consequence of the modest expansion of the Spemann organizer during their gastrula stage of development.

## 2. Results

### 2.1. Spontaneously variant females: F0 (founder) and F1 generations

Three spontaneously variant females (designated F0-1 to −3, Fig. 1A) were identified in conventional crosses with commercially available farmed *Xenopus laevis*. A large proportion of their offspring exhibited a range of enhanced dorsoanterior phenotypes, independent of the mating male. To facilitate our analysis, we categorized the variant larvae into three classes based on the extent of dorsoanteriorization at stage 28/29 (panels and table in Fig. 1A): Class I with enlarged head, laterally extending cement gland, swollen trunk, and shortened tail (see also Fig. 2A, B); Class II with peripherally extending cement gland, reduced trunk, and rudimentary tail; Class III with cement gland surrounding body and trunkless, tailless, and cylindrically symmetric body.

**Fig. 2.**
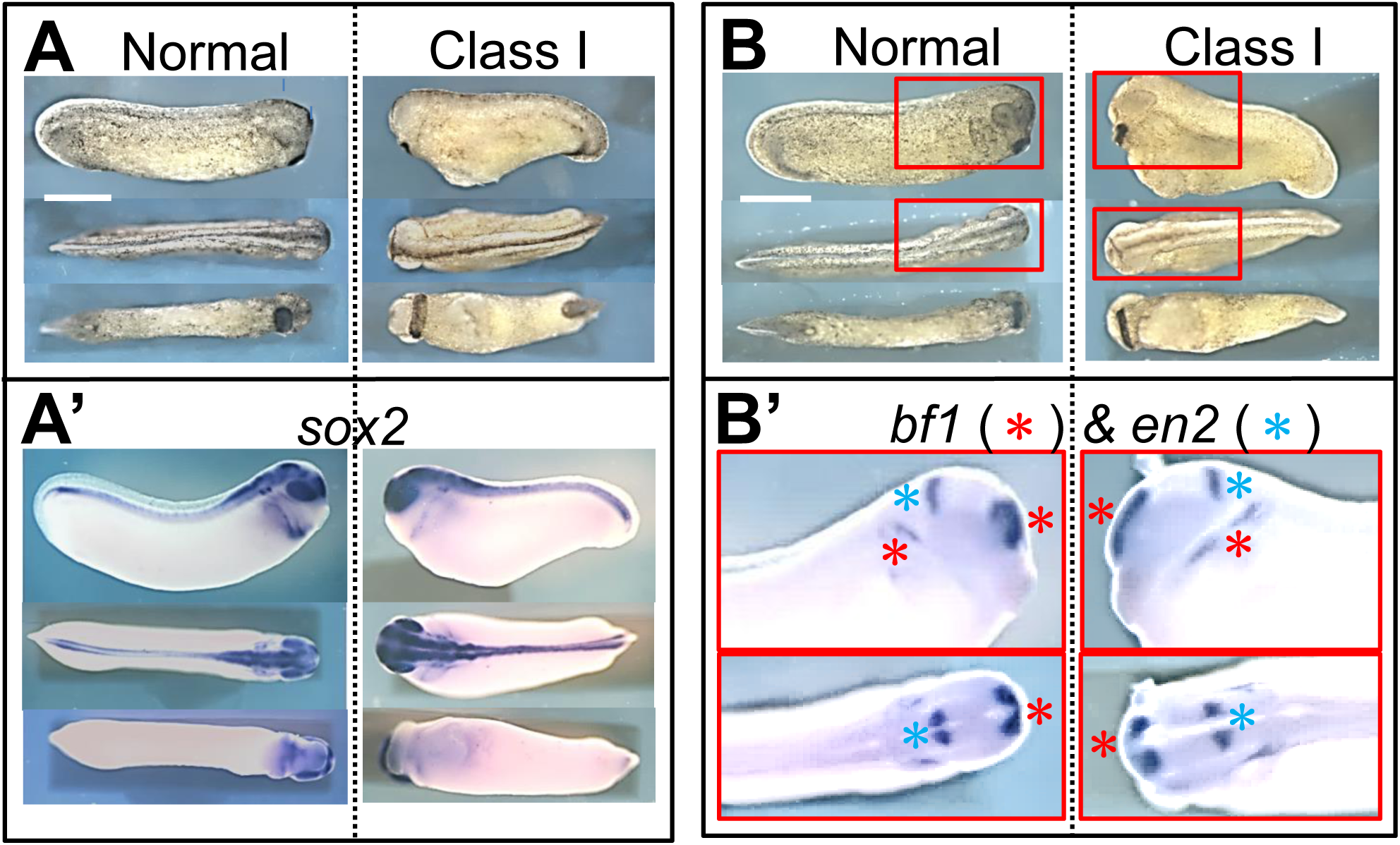
Enlargement of brain structures in Class I larvae. (A, B) Morphology of normal and Class I larvae at stage 29/30 used for WISH analysis in (A’) and (B’). The top, middle, and bottom panels in (A) and (B) show lateral, dorsal, and ventral views, respectively. Scale bar: 1 mm. (A’) Expression of *sox2* in normal and Class I larvae. The arrangement of the images is the same as in (A). (B’) Expression of *bf1* (red asterisk) and *en2* (light blue asterisk) in normal and Class I larvae. Red boxed areas in (B) are magnified.

F0-1 larvae with normal morphology were reared to sexual maturity (F1 generation) to assess whether the propensity of the F0 generation to produce larvae with enhanced dorsoanterior phenotypes was transmitted to the subsequent F1 generation. For this purpose, 5-7 randomly selected F1 females were artificially inseminated with sperm from a single normal male. In each of five such groups, the majority of F1 females produced offspring with variant phenotypes, albeit to varying degrees, whereas F1 males did not produce variant offspring when mated to normal females, confirming the primary role of maternal effects on variant phenotypes. Data from a total of 28 F1 females are summarized in Fig. 1B. They show that the propensity of F0-1 to produce the variant offspring is fairly well transmitted to the females of the F1 generation, suggesting that the enhanced dorsoanterior phenotypes are due to the genetic mutation, although the phenotypic penetrance appears to be incomplete. In the second outcross, variant females did not necessarily reproduce the same proportion of variant phenotypes as in the first cross. Similar non-reproducibility was consistently observed in subsequent outcrosses. The reasons for the variability between sibling females (Fig. 1B) and even at the individual level remain unknown, but we noted that the production of variant offspring tended to increase with maternal age and mating interval. In the following experiments, offspring of F0 and F1 females were collected and fixed at the desired developmental stage and analyzed only if the sibling of the offspring showed a high frequency of variant phenotypes at stage 29/30.

### 2.2. Enlargement of brain structures with rostral bias in Class I larvae

To examine brain structures within the enlarged head of Class I larvae, we first visualized the expression of *sox2*, a pan-neural marker, at stage 29/30 by whole-mount *in situ* hybridization (WISH) (Fig. 2). The rostral part of the brain and the eyes appeared to be significantly enlarged in Class I larvae compared to normal larvae (Fig. 2A’). Further examination at the level of brain subdivisions revealed expanded expression of *bf1*, a specific marker of the forebrain and anterolateral placode, whereas *en2*, which is expressed specifically at the midbrain-hindbrain boundary, appeared less affected (Fig. 2B’). Similar WISH analyses of the other three Class I tailbuds yielded consistent results, confirming the rostral bias of brain enlargement in Class I tailbuds.

Brain enlargement was further investigated at the cellular level by examining nuclear stained serial cross-sections of the Class I tailbud at stage 29/30. Each 6th section and contiguous series of sections are shown in Fig. 3 and Fig. S1, respectively. Comparison with the control revealed significantly increased cell populations in the forebrain (Fig. 3, sections s7-s25) and forebrain-derived eyes (sections s31-s43), whereas those in the hindbrain remained relatively unchanged (section s78). Assessment of midbrain cell populations was difficult because the rostral part of the neural tube bends ventrally during the relevant developmental period (Hausen and Riebesell, 1991), as confirmed by *bf1* expression in Fig. 2B’. Consequently, forebrain and midbrain regions appeared on the same cross-section at certain points (s31-s37 of normal and s31-s43 of variant brain sections). DAPI nuclear staining could not identify the two regions separately, so the corresponding regions on these sections are labeled ‘1/2’ (‘forebrain/midbrain’). Note that the apparent ventral extension of the forebrain actually represents its rostral extension in terms of neural tube structure. Essentially the same results were obtained for the other two sets of experiments, with some variation in the degree of enlargement and the range of enlarged rostral parts in the variant brain (individual data not shown).

**Fig. 3.**
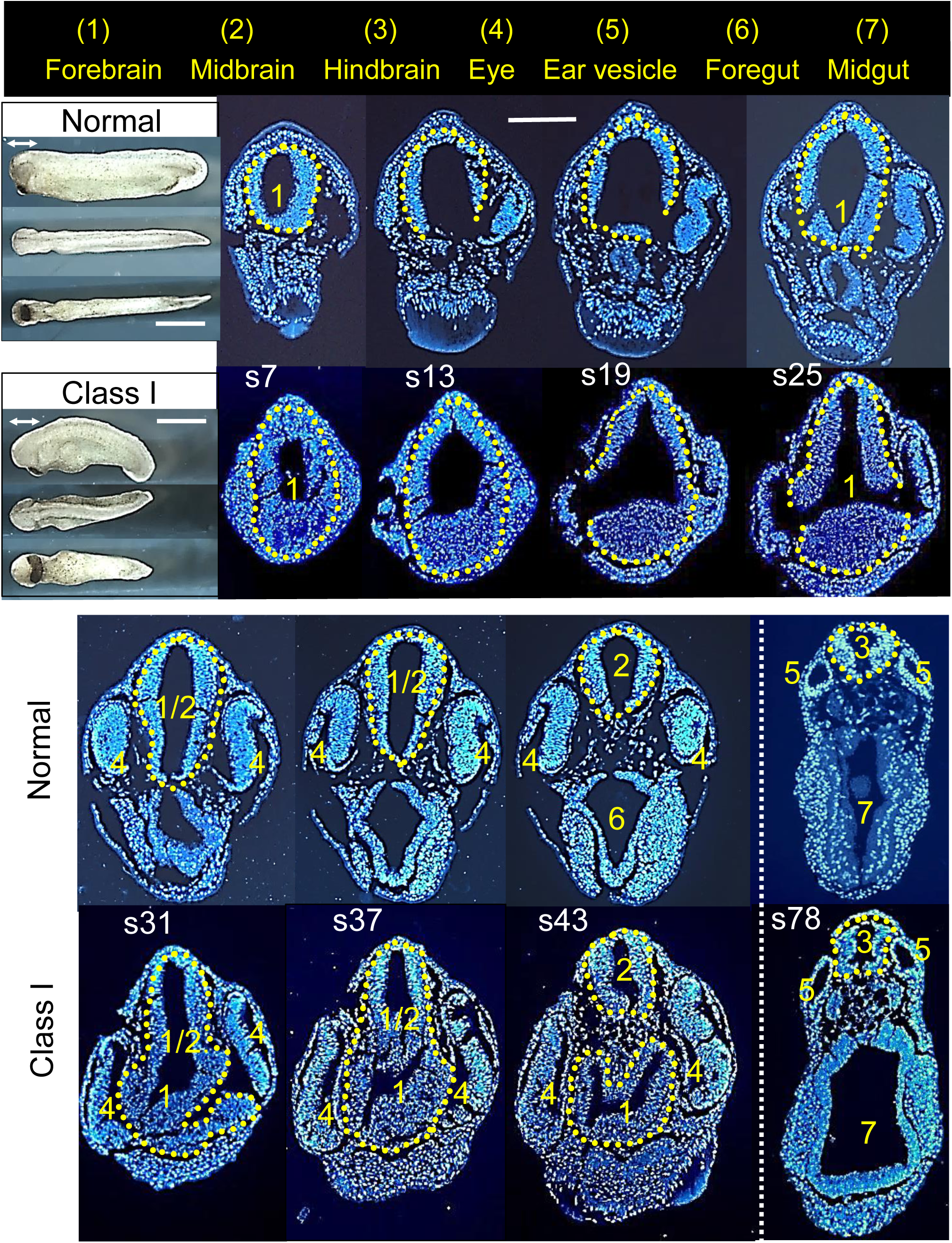
Serial cross-sections through the head of normal and Class I tailbud larvae at stage 29/30. Serial cross-sections were taken from normal and Class I larvae shown in bright field at upper left with scale bar: 1 mm. The part of the head indicated by the bidirectional arrow in the lateral view was sectioned at 8μm and the resulting serial sections were stained with DAPI. Every 6th section from section 7 (s7) to section 43 (s43) and a typical section at some distance (s78) are shown for normal (upper row) and Class I (lower row) larvae. The tissues of interest in the sections are indicated by the numbers given to each tissue at the top of Fig. 3. Specifically, forebrain (1), midbrain (2), and hindbrain (3) are outlined with yellow dotted lines. Scale bar: 0.25mm.

To follow brain development in Class I larvae, serial cross-sections were taken from tadpoles corresponding to stage 40 and stained with hematoxylin/eosin to examine changes in brain tissue architecture. Results for the most severely affected Class I tadpole are shown in Fig. 4 for each 8th section and in Fig. S2 for contiguous series of sections. Compared to a normal stage 40 tadpole, this Class I tadpole showed marked enlargement of the forebrain, midbrain, eyes, and to a lesser extent the spinal cord. The degree of increase in both neural cell population and ventricular size within these neural tissues was more pronounced toward the rostral side. This synchrony between the two indices was expected because the inner surface of the ventricles is composed of neuroepithelial cells, which are precursors of neuronal and glial cells. Analysis of another stage 40 equivalent Class I tadpole, which is less severely affected in appearance than the specimen analyzed in Fig. 4, showed that the significant enlargement was mainly confined to the forebrain region. Taken together, the brains of the Class I variants show a rostral-biased enlargement across the larval period from stage 29/30 tailbuds to stage 40 tadpoles, although the degree of enlargement and the area affected are not necessarily invariant among individuals.

**Fig. 4.**
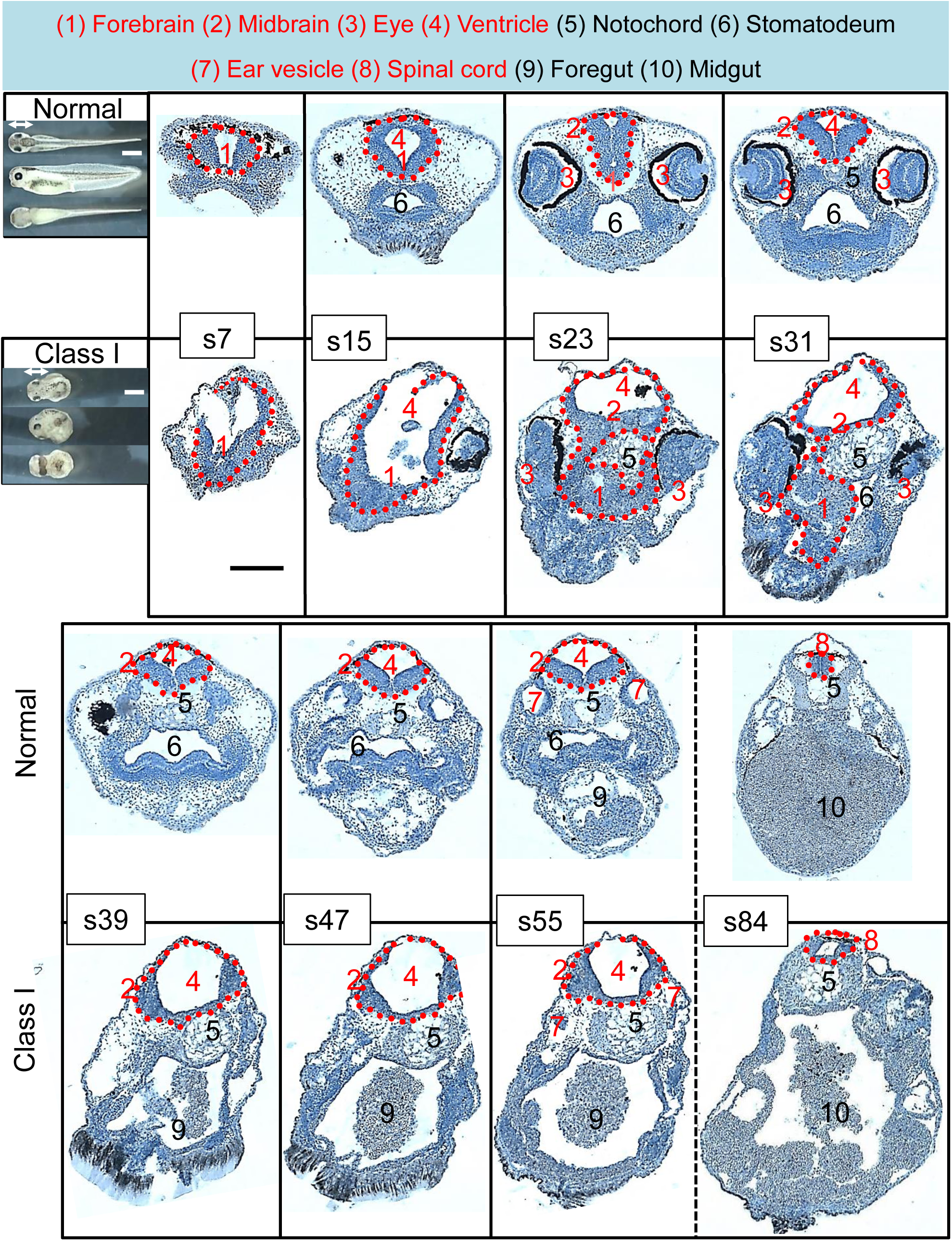
Serial cross-sections through the head of normal and Class I tadpoles at stage 40. Serial cross-sections were taken from normal and Class I larvae shown in bright field at upper left with scale bar: 1 mm. The part of the head indicated by the bidirectional arrow in the dorsal view was cross-sectioned at 15μm and the resulting serial sections were stained with H/E. The tissues of interest in the sections are indicated by the numbers assigned to each tissue at the top of Fig. 4. Neural tissues are indicated by a red number. Specifically, forebrain (1), midbrain (2), and spinal cord (8) are outlined with red dotted lines. Scale bar: 0.25 mm.

### 2.3. Rostral-biased enlargement of neural plate structures in neurulae of variant females

We investigated the onset of brain enlargement in variant offspring by examining *sox2* expression. WISH analysis revealed its lateral expansion in many of the stage 13 neurulae of variant females, indicating lateral expansion of the neural plate region, i.e., the brain primordium (Fig. 5A). This and subsequent experiments with stage 13 and pre-stage 13 embryos were performed on unselected synchronous embryos grown from artificially fertilized eggs, as there is little visible difference between normal and variant embryos up to stage 13. To quantify *sox2* expression, we measured the width of the *sox2*-positive region (d) relative to that of the embryo body (D), as illustrated in Fig. 5A. The normalized d/D values, plotted in histograms, confirmed that *sox2* expression was laterally expanded in the majority of F0-1 neurulae (Fig. 5A’), albeit with considerable inter-individual variability in this expansion. The expansion of the neural plate should contribute to future brain enlargement, as confirmed in Figs. 2 to 4, since a large proportion of *sox2*-positive cells located in the central part of the neural plate give rise to neurogenic cells that populate the inner surface of the brain ventricles (neuroepithelium) during neural tube formation. The remaining *sox2*-positive cells in the periphery are neural crest cells, the precursors of various peripheral derivatives.

**Fig. 5.**
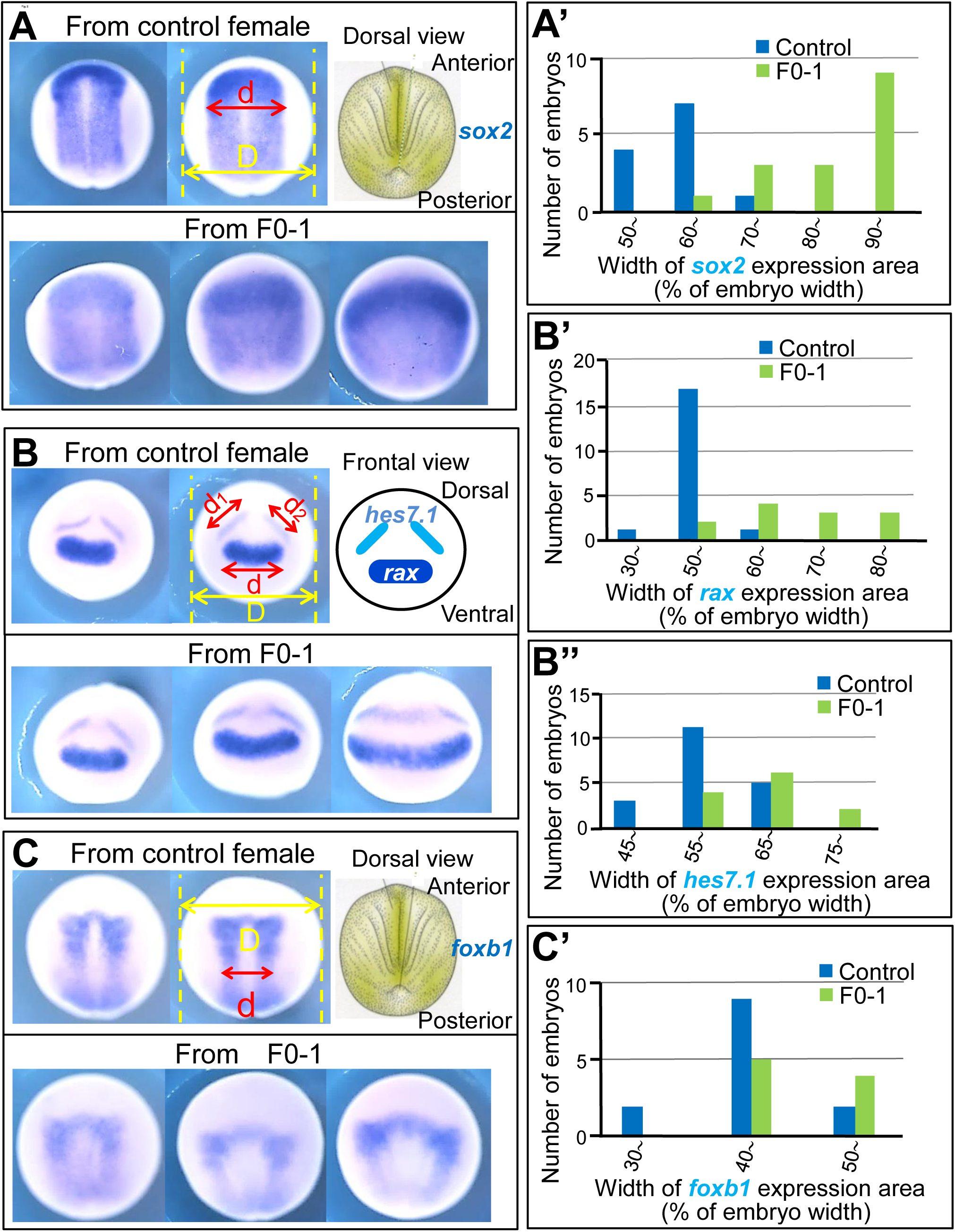
Enlargement of the neural plate and its subdivisions in stage 13 neurulae of variant females. (A, B, C) Examples of *sox2* (A), *rax* and *hes7.1* (B), and *foxb1* (C) expression in stage 13 neurulae from control female (upper panels) and from F0-1 (lower panels). (A’) Histograms generated from *sox2* expression data. The abscissa is the d/D value (% of embryo width). The mean ± SD for control and F0-1 neurulae are 62.6 ± 3.8 and 86.2 ± 10.2, respectively. The difference between the two groups is significant by t-test at p =1.6 × 10 ^−14^. (B’, B’’) Histograms generated from *rax* (B’) and *hes7.1* (B’’) expression data. Abscissa is d/D in (B’) and (d_1_+d_2_)/D in (B’’). Mean ± SD values (% of embryo width) for control and F0-1 neurulae are 54.4 ± 6.2 and 71.0 ±12.3 in *rax*, and 61.0 ± 6.1 and 69.0 ± 10.9 in *hes7.1* expression, respectively. Differences between the two groups ^are significant by t-test at p^=3.0 × 10^−11^ for *rax* expression and p=8.5 × 10^−7^ for *hes7.1* expression. (C’) Histograms generated from *foxb1* expression data. The abscissa is the d/D value (% of embryo width). The mean ± SD for control and F0-1 neurulae are 43.2 ± 6.0 and 46.8 ± 4.7, respectively. The difference between the two groups is significant by t-test at p=1.3 × 10^−3^.

We then examined the enlargement of neurogenic subdivisions in the neural plate using sibling stage 13 neurulae. WISH analysis was performed to assess the expression of *rax*, a specific marker of the future forebrain including the prospective ocular region at stage 13 (Fig. 5B, ‘d’), the specific expression of *hes7.1* at the future midbrain/hindbrain boundary (Fig. 5B, ‘d_1_’ and ‘d_2_’), and the expression of *foxb1* in the future posterior hindbrain region at the level of the prospective medulla region (Fig. 5C, ‘d’; Eagleson and Harris, 1990). For quantitative comparison, we measured the width of expression of each marker gene relative to that of the embryo body, as shown in Fig. 5B and Fig. 5C, and constructed histograms of normalized values for the expression of each gene as for *sox2* (Figs. 5B’, B’’, and C’). The data showed that the expression of all three genes was expanded in many of the F0-1 neurulae, but the degree of expansion appeared to differ between these genes. To confirm this difference, we defined the deviation of expression of each gene in variant neurulae from the normal control (Δ) as [(mean for F0-1 neurulae – mean for control neurulae) / mean for control neurulae]. The Δ for *rax*, *hes7.1*, and *foxb1* were 0.31, 0.13, and 0.08, respectively, indicating a rostral bias in the degree of expansion between the three neural plate subdivisions examined. Essentially the same results were obtained in the other three sets of experiments, but a slight variation was observed, with a tendency for the values of deviation in the expression of *hes7.1* and *foxb1* to be smaller relative to the values for *rax*. This may explain the variation in the extent of rostral bias that we observed in stage 29/30 and 40 variant larvae. Taken together, these results suggest that the rostral-biased brain enlargement in Class I larvae originates as early as the formation of the brain primordium.

### 2.4. Ventral expansion of the Spemann organizer in gastrulae of variant females

In *Xenopus laevis*, neural plate formation is initiated by anti-BMP signaling proteins such as Chordin, Noggin, and Cerberus, which are secreted by the Spemann organizer and act as rostral neural inducers (Bouwmeester et al., 1996; Harland and Gerhart, 1997; Sasai et al., 1994; Stern, 2005). To investigate potential changes in the structure and function of the organizer tissue in the offspring of variant females, we analyzed the expression pattern of *chordin* and *cerberus* in stage 11 gastrulae using WISH. Representative results are shown in Fig. 6A, A’ for *chordin* and Fig. 6B, B’ for *cerberus*. Expression of both genes appeared to be expanded ventrally in variant gastrulae (Fig. 6A, B). To quantify *chordin* expansion, the angle of the *chordin*-expressing region in the border area around the circular blastopore was measured from the center of the blastopore (‘Angle’ in Fig. 6A, top right panel). The cumulative histograms in Fig. 6A’ show that a large proportion of stage 11 gastrulae from F0-1 exhibited a pronounced ventral expansion of *chordin* expression, indicating a substantial expansion of the Spemann organizer tissue toward the ventral side. This was corroborated by quantitative analysis of *cerberus* expression in the same batch of gastrulae using the values of ‘d’ for *cerberus* expression and ‘D’ for embryonic body size (Fig. 6B’), with each term defined in Fig. 6B, top middle panel. Using stage 11 gastrulae from other variant females (F0-2, −3, and F1), the essentially similar pattern of Spemann organizer expansion was observed; the females used here were confirmed to produce a high frequency of stage 29/30 tailbud variant siblings. The considerable variability in the extent of expansion in the variant population, as exemplified in Fig. 6A’ and 6B’, may account for the phenotypic diversity observed in the larvae (Fig. 1A, Class I-III) and neurulae (Fig. 5) of variant females. The rostral-biased brain enlargement in larvae with a relatively mild phenotype (Class I) is thought to be the result of a moderate expansion of the Spemann organizer during the gastrula stage.

**Fig. 6.**
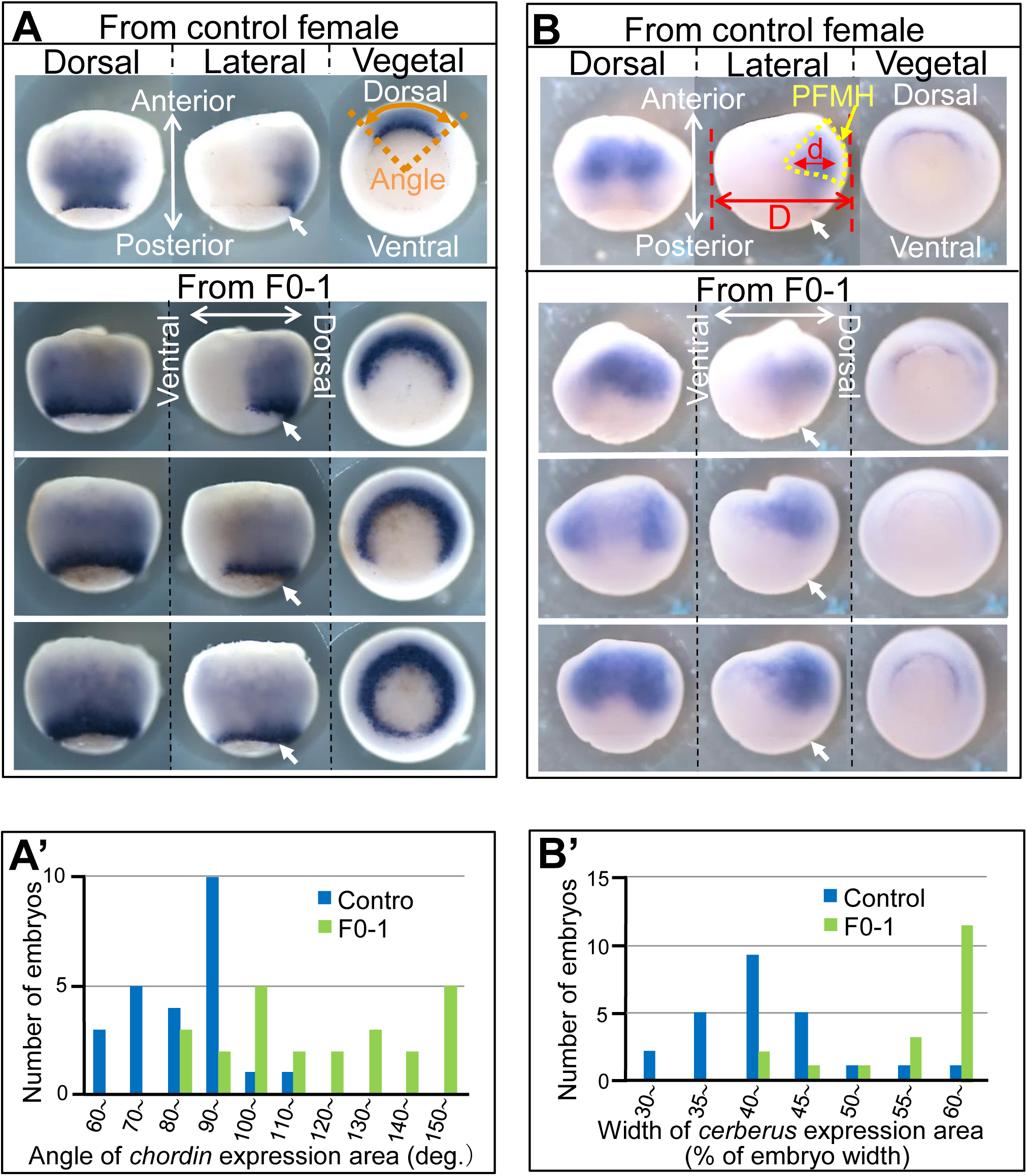
Expanded expression of *chordin* and *cerberus* in gastrulae of variant females. (A, B) Examples of *chordin* (A) and *cerberus* (B) expression in stage 11 gastrulae from control female (top panels) and those from F0-1 (lower panels). Dorsal, lateral, and vegetal views are shown in the left, middle, and right panels, respectively. PFMH in the top middle panel indicates prospective forebrain, midbrain, and hindbrain regions (used for discussion in *3.1)*. (A’) Histograms generated from *chordin* expression data. The abscissa is the value of the expression angle defined by ‘Angle’ in (A). The mean ± SD values (degrees) for control and F0-1 embryos are 86.1 ± 12.2 and 130.6 ± 47.3, respectively. The difference between the two groups is significant by t-test at p=3.0 × 10^−25^. (B’) Histograms generated from *cerberus* expression data. The abscissa is d/D value (% of embryo width). The mean ± SD for control and F0-1 embryos are 43.7 ± 8.3 and 62.4 ± 11.9, respectively. The difference between the two groups is significant by t-test at p _= 5.4 × 10_^−19^.

### 2.5. Ventral expansion of siamois expression in blastulae of variant females

Activation of *chordin* and *cerberus* requires prior expression of *siamois*, a homeodomain transcription factor gene in *Xenopus* (Kessler, 1997). We then examined whether the expression pattern of *siamois* was altered in the offspring of variant females. In stage 9.5 late blastulae of F0-1, *siamois* was expressed dorsally near the equatorial region, similar to the control, but in many cases the expression area expanded laterally toward the ventral side compared to the control (Fig. 7A). Cumulative histograms constructed using the values of ‘d’ for *siamois* expression and ‘D’ for embryonic body size, with each term defined in Fig. 7A, quantitatively confirmed the lateral expansion of *siamois* expression (Fig. 7A’). Using the offspring of F0-2, −3, and F1 females, which were confirmed to produce a high frequency of stage 29/30 tailbud variant siblings, the same pattern of expanded *siamois* expression was observed (e.g., Fig. 7A’’).

**Fig. 7.**
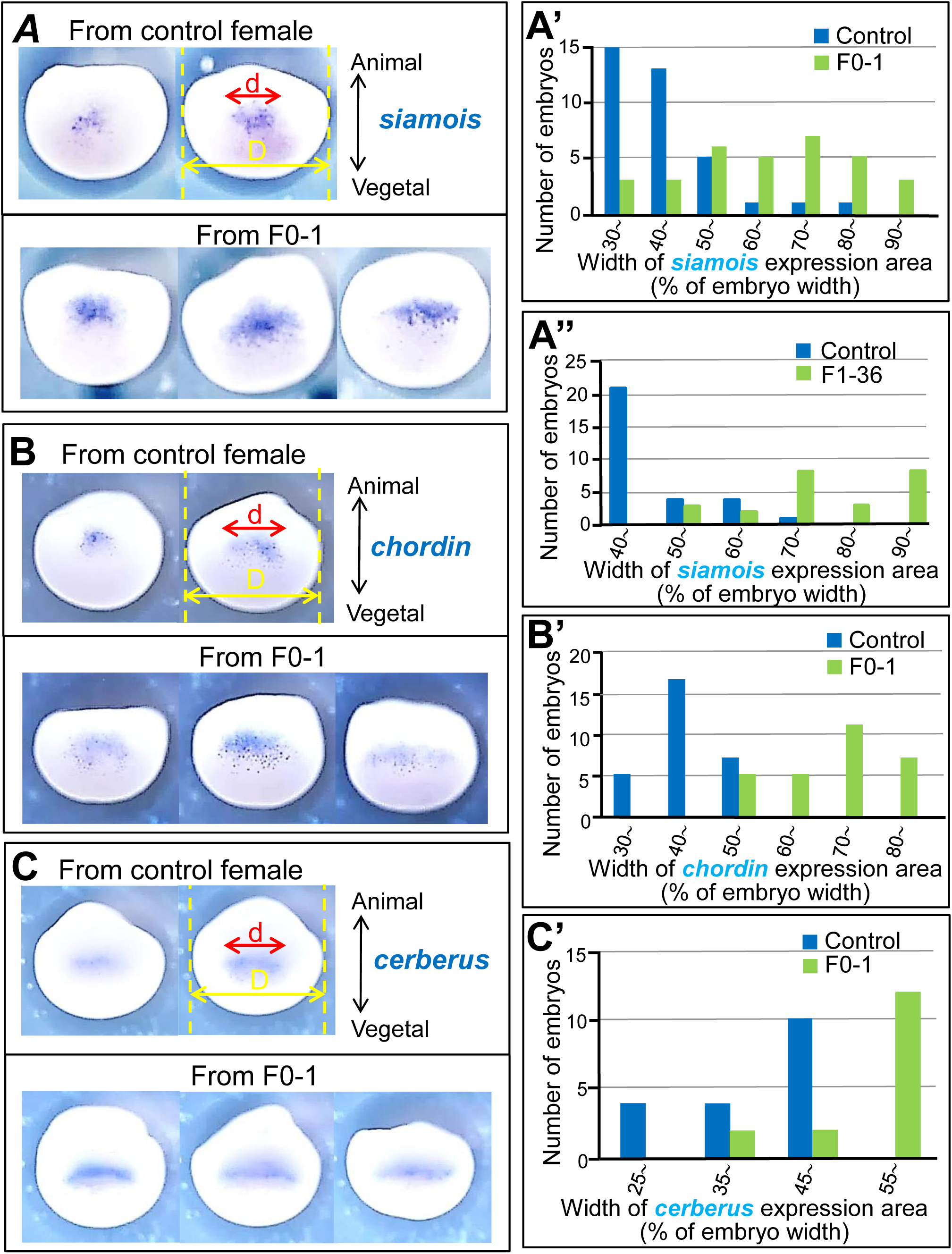
Expanded expression of *siamois* in blastulae of variant females. (A, B, C) Examples of *siamois* (A) and *chordin* (B) expression at stage 9.5 and *cerberus* (C) expression at stage 10 in embryos from control female (upper panels) and F0-1 (lower panels). Dorsal views are shown. (A’, A’’) Histograms generated from *siamois* expression data. Blastulae from F0-1 (A’) and F1-36 (A’’) were used. F1-36 is a progeny of F0-1 from a different cross than the one that produced the F1 progeny listed in Fig. 1B. Abscissas are d/D values (% of embryo width). Mean ± SD values for control and F0-1 embryos are 42.8 ± 11.6 and 64.4 ± 16.8, respectively (A’), and 48.2 ± 8.0 and 77.0 ± 13.2 for control and F1-36 embryos, respectively (A’’). Differences between control and variant groups are significant by t-test at p=5.9 × 10^−32^ for (A’) and p=1.8 × 10^−31^ for (A’’). (B’, C’) Histograms generated from *chordin* (B’) and *cerberus* (C’) expression data. Abscissas in (B’) and (C’) are d/D values (% of embryo width) with each term defined in (B) and (C), respectively. Mean ± SD values for control and F0-1 embryos are 44.4 ± 5.3 and 70.3 ± 10.0 for *chordin* expression, respectively (B’), and 43.3 ± 9.1 and 54.4 ± 7.1 for *cerberus* expression, respectively (C’). The difference between control and F0-1 groups is significant by t-test at p=1.4 × 10^−36^ for *chordin* expression and p=8.8 × 10^−13^ for *cerberus* expression.

We next extended our analysis to the expression patterns of *chordin* and *cerberus* in stage 9.5 late blastula and stage 10 early gastrula, respectively, using the F0-1 sibling used for *siamois* expression analysis. Both genes were expressed in the dorsal equatorial region as in the control, but their expression expanded laterally in the majority of the offspring of the variant female (Fig. 7B, C), resembling the expression pattern of *siamois* (Fig. 7A). The expansion of *chordin* and *cerberus* expression was confirmed quantitatively as in *siamois* (Fig. 7B’, C’). The considerable variability in the expanded expression of *chordin* (Figs. 6A’ and 7B’) and *cerberus* (Figs. 6B’ and 7C’) in the variant population could be attributed to a similar variability in the expansion of the preceding *siamois* expression (Fig. 7A’, A’’). Collectively, these results indicate that *siamois*-expressing cells are distributed in a ventrally spreading manner in variant blastulae, leading to a ventral expansion of the Spemann organizer tissue during the gastrula stage.

### 2.6. Dispersed distribution of maternal wnt11b mRNA in the vegetal cortical cytoplasm (VCC) of variant female eggs

The expression of *siamois* in dorsal cells requires the prior formation of the nuclear β-catenin/Tcf3 transcription complex in *Xenopus* (Brannon et al., 1997). Cytoplasmic β-catenin is stabilized by Wnt/β-catenin signaling to enter the dorsal cell nucleus and form the complex. Thus, the ventrally expanded distribution of *siamois*-expressing cells reflects a prior expansion of the active domains of the Wnt/β-catenin pathway in a similar pattern. Notably, this pathway has been shown to be activated in the dorsal embryonic region by maternally derived dorsalizing factors. They initially localize to the VCC in newly laid eggs and are translocated dorsally following fertilization by the microtubule-mediated transport systems, including cortical rotation that moves the entire VCC dorsally along an animal-vegetal meridian (Gerhart, 2004; Houston, 2012, 2017; Weaver and Kimelman, 2004). Since maternal *wnt11b* mRNA is a promising candidate as such a dorsalizing factor (Tao et al., 2005), it is conceivable that the expansion of active regions of Wnt/β-catenin signaling is due to an increased level and/or distribution of maternal *wnt11b* mRNA in the eggs of variant females. To confirm this possibility, we analyzed mRNA levels by RT-PCR in unfertilized eggs laid by 15 of the 28 F1 females listed in Fig. 1B (Fig. 8A). These mothers produced offspring with varying degrees of dorsoanterior phenotypes (Fig. 1B). The quantified data show that wnt11b mRNA levels (Fig. 1B, solid circles) do not correlate with the degree of dorsoanteriorization in the offspring of the 15 mothers examined (Fig. 1B, percentage bars). This indicates that differences in the abundance of maternal wnt11b mRNA deposited in the laid eggs are not the primary determinant of the different dorsoanterior phenotypes in the developing offspring of the variant females.

**Fig. 8.**
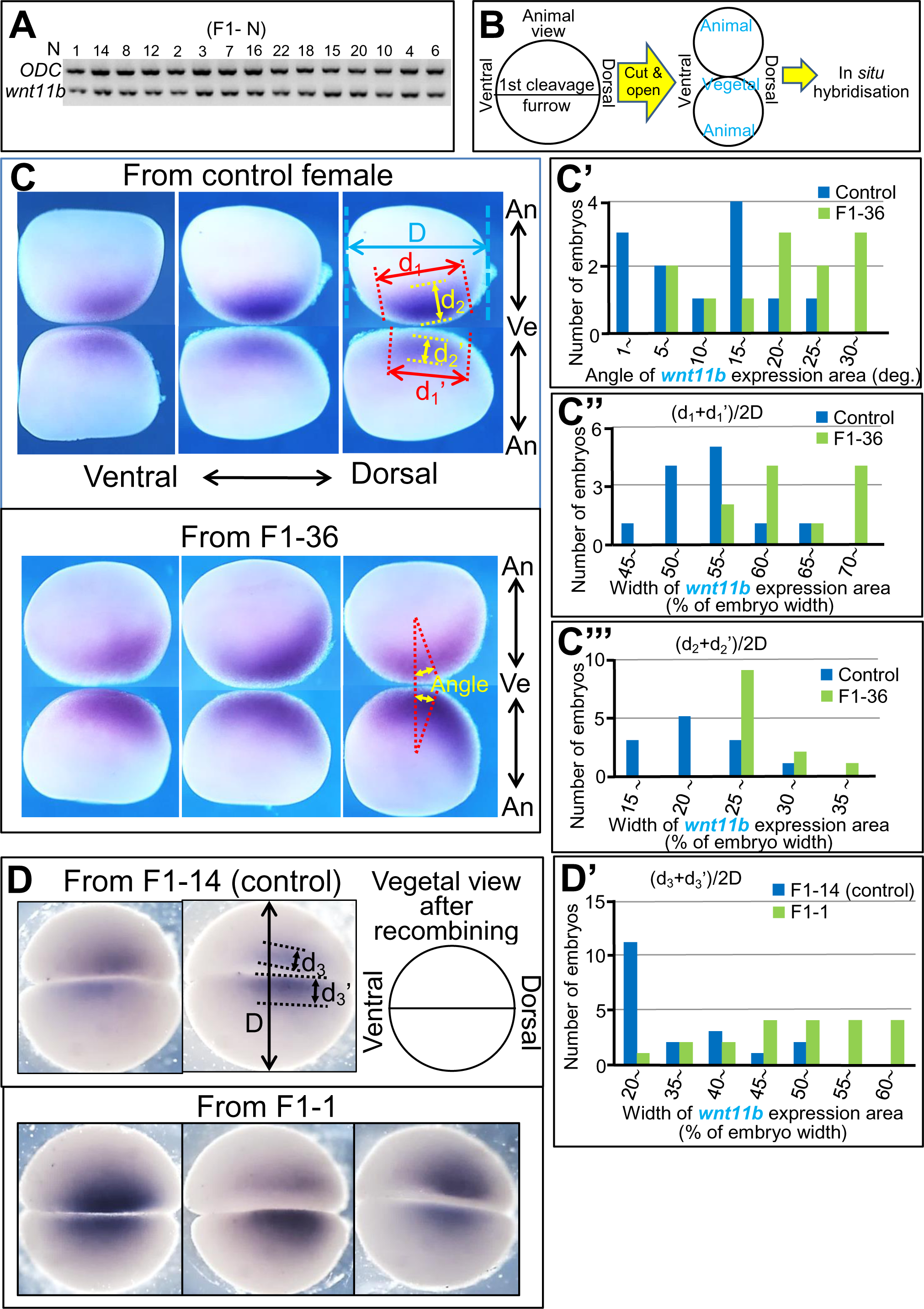
Dispersion of maternal *wnt11b* mRNA in the vegetal cortical cytoplasm of variant female eggs. (A) RT-PCR products of *wnt11b* mRNA in unfertilized eggs of females of the F1 generation. N in (F1-N) indicates the assigned number of F1 females examined as shown in Fig. 1B. (B) Protocol for WISH with the bisection procedure and method for displaying stained samples in (C). (C, D) Examples of *wnt11b* mRNA expression at stage 2^−^. In (C), cut-surfaces of WISH-processed embryos from control female (upper panels) and F1-36 (lower panels) are shown in the arrangement illustrated in (B). In (D), embryos from control F1-14 (upper panels) and variant F1-1 (lower panels) were used. Vegetal views of the recombined embryos are shown. (C’) Histograms generated from the rotational shift data of *wnt11b* mRNA. The abscissa is the mean of the angles defined by ‘Angle’ on the pair of cut surfaces in (C), lower right panel. The mean ± SD (degrees) for control and F1-36 embryos are 12.8 ± 7.7 and 22.0 ± 9.7, respectively. The difference between the two groups is significant by t-test at p =1.3 × 10^−7^. (C’’, C’’’, D’) Histograms generated from the dispersion data of *wnt11b* mRNA. Abscissas are (d_1_ + d_1_’)/2D in (C’’), (d_2_ + d_2_’)/2D in (C’’’), and (d_3_ + d_3_’)/2D in (D’), respectively, with each term defined in (C), upper right panel, or (D), upper panel. Mean ± SD values (% of embryo width) for control and variant F1 embryos are 56.3 ± 5.2 and 66.7 ± 5.8 for dorsoventral dispersion in (C’’), 23.1 ± 4.7 and 29.1 ± 2.9 for inward dispersion in (C’’’), and 34.7 ±9.3 and 59.0 ± 9.9 for lateral dispersion in (D’), respectively. Differences between control and variant F1 groups are significant by t-test at p=2.7 × 10^−10^ in (C’’), p=2.2 × 10^−7^ in (C’’’), and p=1.2 × 10^−17^ in (D’).

We then examined the distribution pattern of maternal *wnt11b* mRNA in stage 2^−^embryos of variant and control females, identified by the appearance of the first cleavage furrow after fertilization. They were fixed and sectioned sagittally along the future first cleavage plane to facilitate WISH analysis (Fig. 8B). As expected for a dorsalizing factor, *wnt11b* mRNA was distributed in the VCC and appeared to shift from the animal-vegetal pole axis to the dorsal direction in stage 2^−^embryos of both variant and control females (Fig. 8C). For quantitative comparison, we measured the angle of rotation of the *wnt11b* mRNA-positive region (‘Angle’ in Fig. 8C, lower right panel), which was 22.0 + 9.7 degrees and 12.8 + 7.7 degrees in embryos from variant and control females, respectively (Fig. 8C’). The significance of these different values is uncertain because cortical rotation is a highly variable process with a range of 13-35 degrees, although compatible with normal development (Vincent and Gerhart, 1987). Nevertheless, it can at least be said that cortical rotation is not significantly affected in fertilized eggs of variant females.

Another interesting point in Fig. 8C is that *wnt11b* mRNA appears to be more diffusely distributed in the VCC compared to controls. To quantify this dispersed distribution, we measured the width of the *wnt11b* mRNA-positive region (d_1_ and d_1_’) relative to that of the embryo body (D), as shown in Fig. 8C, upper right panel. Histograms of normalized (d_1_ + d_1_’)/2D values confirmed that *wnt11b* mRNA was more dispersed along the d_1_ measurement axis in many of the embryos from the variant female compared to control embryos (Fig. 8C’’). When measuring the degree of inward dispersion (d_2_ and d_2_’ in Fig. 8C, upper right panel), histograms of normalized (d_2_ + d_2_’)/2D values showed that *wnt11b* mRNA was also more dispersed inwards in variant embryos (Fig. 8C’’’). Furthermore, a similar analysis of lateral distribution ((d_3_ + d_3_’)/2D, with each term defined in Fig. 8D, upper panel) showed a more dispersed lateral distribution of *wnt11b* mRNA in variant embryos (Fig. 8D’). Taken together, these results indicate that *wnt11b* mRNA is dispersed almost omnidirectionally in stage 2^−^variant embryos. Lateral spreading is of particular interest because it may allow lateral expansion of the active domains of Wnt/β-catenin signaling, leading to the lateral (dorsal to ventral at blastula stage) expansion of *siamois* expression. The variability in the degree of lateral spreading of *wnt11b* mRNA in the variant population (Fig. 8D’) could explain the variability in the ventral expansion of *siamois* expression in the variant blastulae (Fig. 7A’, A’’). These experiments were repeated with stage 2^−^embryos from the other two variant females with essentially the same results.

## 3. Discussion

### 3.1. Enlargement of brain structures due to Spemann organizer expansion

This study is the first to describe a naturally occurring female variant line within a farmed *Xenopus laevis* population that produces offspring with varying degrees of enhanced dorsoventral phenotypes due to maternal effects (Fig. 1, Class I-III). Detailed analysis of the variant offspring with the mild phenotype (Class I) revealed that they have an enlarged brain with a rostral bias during the larval stages, while maintaining the overall basic tissue architecture (Figs. 2 to 4). The characteristic brain enlargement can be traced back to stage 13 neurula (Fig. 5), when the Spemann organizer has induced and begun to pattern the neural plate, i.e., the brain primordium. Since we have also shown that the Spemann organizer is expanded ventrally to varying degrees in variant offspring (Fig. 6), it is reasonable to assume that the brain enlargement observed in Class I larvae is a consequence of a modest expansion of the Spemann organizer. Several lines of experimental evidence support this notion of a causal relationship between organizer tissue size and brain morphology, as discussed below.

A range of enhanced dorsoanterior phenotypes similar to those observed in this study (Fig. 1A), including Class I-like phenotypes as inferred from their external appearance, have been produced by exposing 32-to 128-cell stage embryos to 0.3 M LiCl for 10-12 min (Kao and Elinson, 1988) or by exposing fertilized eggs to 30-70% D_2_O for a few minutes during the first third of the first cleavage (Scharf et al., 1989). Li treatment broadly activates the Wnt/β-catenin pathway in the embryo by inhibiting GSK-3β activity in the pathway, leading to enhanced dorsalization (Hedgepeth et al., 1997; Klein and Melton, 1996), whereas D_2_O treatment randamizes the normally parallel microtubule assembly, which has been suggested to cause ventral expansion of the dorsalizing factors (Scharf et al., 1989). In both cases, the resulting enhancement of dorsal and anterior structures has been attributed to an increase of Spemann organizer tissue. Indeed, our own experiments showed that D_2_O treatment of fertilized eggs from normal parents caused a ventral expansion of *chordin* expression similar to that observed in the offspring of the variant female (Fig. S3A, compare with Fig. 6A’), confirming the ventral expansion of the Spemann organizer by D_2_O treatment. In addition, a large proportion of D_2_O-treated neurulae showed expanded expression of the pan-neural marker *sox2* (Fig. S3B, C) and rostral neural markers such as *bf1* (Fig. S3B, C), *emx1,* and *fez* (Fig. S3C).

Experiments using morpholino oligomers (MOs) to manipulate gene expression have also supported a causal link between organizer size and brain morphology. Injection of ONT1-MO (Inomata et al., 2008) or ADMP-MO (Reversade and de Robertis, 2005) at the 4-cell stage induced a Class I-like phenotype as judged by external appearance. In MO-injected gastrulae, the organizer region was shown to be expanded ventrally. This is expected because ONT1 and ADMP normally act as fine-tuning ventralizing factors in the dorsal embryonic region by increasing BMP signaling, and thus the MOs against them are predicted to counteract their ventralizing BMP activity, leading to some modest expansion of the organizer tissue. Taken together, these previous and our present results suggest that brain enlargement in Class I larvae is most likely due to a modest expansion of the Spemann organizer that could result from a limited number of genetic mutations.

The expression domains of *chordin* and *cerberus* nearly overlap, but *cerberus* appears to be expressed slightly anterior to chordin, primarily in the anterior endomesodermal region (compare Fig. 6A and 6B), as previously reported (Koide et al., 2002). In particular, *cerberus* expression does not extend to the dorsal lip of the blastopore (Bouwmeester et al., 1996), whereas *chordin* expression does (Fig. 6A and 6B, white arrows). Thus, *cerberus*-expressing cells mainly underlie the prospective forebrain, midbrain and hindbrain regions of the gastrula dorsal ectoderm (labeled PFMH in Fig. 6B; Keller et al., 1992), suggesting a primary role for Cerberus in the development of rostral brain structures. Secreted Cerberus, together with Chordin, is thought to prevent differentiation of the PFMH region into epidermal tissue by sequestering ectoderm-derived BMP protein from its receptor on PFMH cells (Bouwmeester et al., 1996). Cerberus also appears to inhibit the differentiation of PFMH cells into mesodermal tissue by sequestering endomesoderm-derived Nodal and Wnt proteins (Piccolo et al., 1999). The differentiation of PFMH into neuroectoderm with a rostral neural identity is indicated to be autonomously promoted by intrinsic FGF/Ets signaling in gastrula ectodermal cells (Hongo and Okamoto, 2022). Subsequently, a rostrocaudal gradient of caudalizing factors such as FGF and Wnt secreted by the broad equatorial mesoderm appears to give the induced neuroectoderm an initial rostral-caudal pattern (Doniach et al., 1992; Hongo et al., 1999; Hongo and Okamoto, 2022; Kiecker and Niehrs, 2001; Stern, 2005); note that the dorsal part of the equatorial mesoderm corresponds to the posterior mesodermal part of the Spemann organizer. The expansion of *cerberus*-expressing and *chordin*-expressing cells in the variant gastrulae (Fig. 6) should contribute significantly to the expansion of rostral neural structures at later stages, which was indeed observed (Figs. 2 to 5).

Of note is the heterogeneous nature of the enlargement of brain subdivisions in the variant offspring, which is more intense rostrally. The pattern of rostral bias was already evident at the stage of neural plate formation (Fig. 5). The most plausible explanation for the origin of this characteristic pattern is that it reflects to some extent the independent modes of action of the multiple neural inducing and patterning factors involved in neural plate formation: for example, if the overall activity of caudalizing factors from the broad equatorial mesoderm is not expanded as effectively as the activity of rostral inducing factors in the variants, then this could result in a rostral bias in the patterning of neural plate subdivisions. Indeed, our preliminary experiments suggest that the pattern of FGF signaling activity in the equatorial mesodermal region as a whole does not appear to be markedly altered in variant gastrulae, as judged by the immunostaining pattern for P-MAPK, an indicator of FGF signaling activity, even though the dorsal portion of the equatorial mesoderm corresponds to the posterior mesodermal portion of the Spemann organizer.

### 3.2. Role of dorsal determinant spreading and subsequent expansion of Wnt/β-catenin signaling in Spemann organizer expansion

Activation of Wnt/β-catenin signaling in the dorsal embryonic region is triggered by dorsalizing factors of maternal origin, commonly referred to as dorsal determinants (Weaver and Kimelman, 2004). Several potential dorsal determinants have been proposed, including Dishevelled (Miller et al., 1999; Weaver et al., 2003), phospho-Lrp6 (Dobrowolski and de Robertis, 2012), Huluwa (Yan et al., 2018), and Wnt11b (Tao et al., 2005). The mRNAs for these proteins have been shown to be presynthesized during oocyte maturation and localized in the cortical cytoplasm adjacent to the vegetal pole of the fully mature oocyte. In the fertilized egg, these mRNAs, and presumably their translation products, are distributed in the egg VCC and migrate from the original vegetal site to the future dorsal equatorial region of the embryo. There they stabilize cytoplasmic β-catenin by activating the Wnt/β-catenin signaling pathway, not necessarily from its upstream end, leading to the activation of *siamois* later in the blastula stage, which is a prerequisite for Spemann organizer formation in *Xenopus*. The migration of dorsal determinants has been shown to be driven either directly by the microtubule-mediated transport system per se, which assembles within the VCC shortly after fertilization (Weaver and Kimelman, 2004), or rather indirectly by cortical rotation, which involves the dorsal directed movement of the entire VCC while retaining dorsal determinants. Notably, cortical rotation itself is also driven by the microtubule-mediated transport system (Gerhart, 2004; Houston, 2012, 2017). In these contexts, two non-exclusive modes of action for Wnt11b have been proposed. First, dorsally transported Wnt11b directly activates the Wnt/β-catenin pathway by binding to the Wnt receptor, thereby stabilizing β-catenin in dorsal regions (Tao et al., 2005). Second, Wnt11b promotes parallel ventral-dorsal assembly of microtubules within the VCC upon fertilization, thereby facilitating microtubule-dependent transport of dorsal determinants to enhance Wnt/β-catenin signaling in dorsal regions (Houston et al., 2022).

Current analyses remain inconclusive as to whether maternal Wnt11b acts by one or both of the proposed mechanisms, nor how the mode of action of maternal Wnt11b is altered in the variant eggs. Experiments with D_2_O-treated fertilized eggs suggest that the precocious formation of a randomly arranged microtubule network causes the transfer of dorsal determinants to broader dorsal-to-ventral equatorial sectors, resulting in enhanced dorsal and anterior structures (Scharf et al., 1989). The observed concomitant reduction of cortical rotation was also attributed to this aberrant microtubule organization. However, in fertilized eggs from variant females, cortical rotation appeared to be unaffected (Fig. 8C, C’), suggesting that functional microtubule assembly was preserved in these eggs. Indeed, preliminary immunostaining experiments showed no signs of precocious or random microtubule assembly in fertilized eggs from variant females. Although the possibility that aberrant microtubule assembly contributes to the enhanced dorsoanterior phenotype in variant offspring cannot be ruled out, it seems more likely that dorsal determinants are dispersed omnidirectionally in the VCC of variant eggs, as in the case of *wnt11b* mRNA (Fig. 8), and move toward the wider equatorial region via microtubule arrays that are functionally assembled, albeit under the control of somewhat dispersed Wnt11b. The resulting dorsal-to-ventral spreading of maternal *wnt11b* mRNA and other candidate dorsalizing factors would facilitate the expansion of the active domains of the Wnt/β-catenin pathway in a similar pattern, resulting in the ventral expansion of *siamois* expression at the blastula stage, which in turn leads to the ventral expansion of the Spemann organizer at the gastrula stage.

Given that the spreading of dorsal determinants and the consequent expansion of active domains of Wnt/β-catenin signaling occurs in all directions, it may seem puzzling that *siamois*-expressing cells expand predominantly in the ventral direction without significant vertical expansion, especially toward the animal pole (Fig. 7). However, this could be explained by the presence of another signaling pathway required for *siamois* expression that acts in concert with the Wnt/β-catenin pathway but, unlike the latter, is active along the broad dorsal-ventral equatorial region of the blastula. The FGF signaling pathway is a prime candidate for such a pathway, as the FGF pathway is normally activated throughout the equatorial region of the blastula (Schohl and Fagotto, 2002) and does not appear to be markedly affected in the variant offspring as noted in *3.1.* Furthermore, the requirement of FGF signaling for *siamois* expression is confirmed by our experiments with FGF2 MO (Fig. S4).

The present study did not address the cause of *wnt11b* mRNA dispersion in variant eggs. Alterations in some cytoplasmic architectures that reduce VCC viscosity may be one possibility. However, to elucidate the exact molecular mechanisms of *wnt11b* mRNA dispersion, we may need further cytological analyses as well as transcriptomic, whole genome and even epigenetic analyses of variant females and/or their eggs. Nevertheless, the variant eggs described in this study will be valuable for identifying the bona fide dorsal determinants and for studying their detailed mode of action.

### 3.3. Implications for the evolution of the vertebrate brain

The evolutionary trend toward a larger cerebral cortex in vertebrates, especially mammals, has long been recognized and is often referred to as neocorticalization. It is noteworthy, however, that brain enlargement extends beyond the cerebral cortex caudally to regions such as the cerebellum, albeit to a lesser extent (Barton, 2002). An evolutionary assessment of the increase in both mass and neuronal number per representative brain subdivision in primates compared to insectivores, which retain several ‘primitive’ mammalian features, shows that the degree of increase is progressively greater toward the rostral side (neocortex > cerebellum > medulla; see Supplementary Information for quantified data and references).

The current investigation shows an apparently analogous trend of rostral bias in the enlargement of brain subdivisions in the larval offspring of the naturally occurring female variant line of *Xenopus*, and suggests that this is a consequence of the expansion of the Spemann organizer. Our analyses further suggest that mutational events in a cascade of intracellular and intercellular signaling events during early embryogenesis could be responsible for organizer expansion. It may be plausible that the rostral-biased brain enlargement observed evolutionarily in extant vertebrates is due, at least in part, to analogous mutational events in ancestral vertebrate lineages that caused expansion of tissues homologous to the Spemann organizer. The corresponding ancestral tissue in the last common ancestor of vertebrates would be the tissue resembling the embryonic shield in teleosts, and tissues that evolved from this shield-like tissue in extant species may include the hypoblast/anterior visceral endoderm/foregut endoderm and Hensen’s node in birds and mice (Houston, 2017; Stern, 2005: Wilson and Edlund, 2001). The anterior visceral endoderm (AVE) in mouse embryos is of particular interest in the context of the present study because it is the topological equivalent of the anterior endomesodermal portion of the *Xenopus* Spemann organizer (Piccolo et al., 1999), which is known to secrete Cerberus as a rostral neural inducer (Bouwmeester et al., 1996). The mouse AVE also secretes Cerberus-like, a homologue of *Xenopus* Cerberus (Belo et al., 1997) and, like the *Xenopus* Spemann organizer, is thought to require nuclear β-catenin activity for its formation (Morkel et al., 2003). Also analogous to the *Xenopus* Spemann organizer, the AVE appears to play a fundamental role in the formation of the rostral brain (Beddington and Robertson, 1999; Belo et al., 1997; Thomas and Beddington, 1996) and in the establishment of rostrocaudal polarity (Perea-Gómez et al., 2002). Thus, some mutations in the AVE could induce a variety of changes in mouse brain development, possibly including rostral-biased enlargement as in *Xenopus*. However, these issues do not seem to have been thoroughly investigated.

It is worth considering the potential consequences of widespread morphological mutations, such as rostral-biased brain enlargement, which may even be lethal. For these mutant lineages to become established as new evolutionary species, additional mutations would have been required to compensate for the lethal effects of the original mutations. In the case of the Class I variants, they failed to survive beyond the tadpole stage due to inappropriate differentiation of digestive tissues. In *Xenopus*, *cerberus*-expressing cells in the anterior endomesoderm contribute to the development of the foregut, midgut and liver (Bouwmeester et al., 1996). Excessive proliferation of *cerberus*-expressing cells (Fig. 6B, B’) may affect not only the induction of neuroectoderm, but also the formation of digestive tissues. Survival of the Class I variant would require additional mutational events to counterbalance the lethal effects of overproduction of *cerberus*-expressing cells on digestive tissue differentiation. It is noteworthy that in the case of adult populations of wild guppies, the increase in brain size (5-10%) of large-brained lines has been shown to be accompanied by a decrease in gut size (8-20%) (Kotrschal et al., 2012). Of interest in this regard is the well-known ‘expensive-tissue hypothesis’, which postulates a trade-off between brain enlargement and gut reduction during vertebrate evolution: this co-evolution of brain and gut has been proposed to occur between average mammals and primates, and in an accentuated form between humans and their primate relatives (Aiello and Wheeler, 1995). The ‘expensive-tissue hypothesis’ is based primarily on metabolic considerations, whereas our present study provides embryological insights in support of it.

## 4. Materials and methods

### 4.1. Xenopus adults and embryos

Adult female and male *Xenopus laevis* were obtained from Hamamatsu Seibutsu Kyouzai Corporation, Hamamatsu. The methods used to maintain them and to obtain embryos by natural mating were as previously described (Mitani and Okamoto, 1989). For artificial insemination, eggs were laid in 0.5 × MBS and fertilized with a sperm suspension in 0.1 × MBS. This was done at around 15°C, as lower temperatures seemed to be more favorable for obtaining a higher proportion of variant embryos. The fertilized eggs were then reared to the desired stage at 15-24°C in 0.1 × MBS. Embryos were staged according to Nieuwkoop and Farber (1967). Animals were handled in accordance with national guidelines and Gakushuin University guidelines for animal experimentation.

### 4.2. Histological procedures

Specimens were fixed in 2% paraformaldehyde and 1% glutaraldehyde in 50mM phosphate buffer for 5 h. After rinsing with buffer, they were dehydrated in graded ethanol and embedded in paraffin according to the standard method. Sequential cross-sections were cut at 8µm thickness for stage 29/30 tailbud samples and at 15µm thickness for stage 40 tadpole samples. After mounting on glass slides, tailbud specimens were sealed for nuclear staining in 25% glycerol containing 1µg/ml DAPI (4’, 6-diamidino-2-phenylindole, dihydrochloride), 100µg/ml DABCO (1, diazabicyclo[2.2.2]octane), and 25µg/ml *p*-phenylenediamine in 20mM Tris-HCl (pH 8.2). Tadpole samples were stained with hematoxylin/eosin using standard procedures.

### 4.3. Whole-mount in situ hybridization

Whole-mount *in situ* hybridization experiments were performed as previously described (Hongo and Okamoto 2022).

### 4.4. Analysis of wnt11b expression by semi-quantitative RT-PCR

Total RNA was prepared from unfertilized eggs using the Proteinase K method (Sambrook et al. 1989). Semi-quantitative RT-PCR was performed as previously described, except that *odc* was used as an internal control instead of *ef1α* (Hongo and Okamoto 2022). Primer sequences were as follows. For *wnt11b*, F: 5’-ACAAAATGCAAGTGCCACGGAG-3’ and R: 5’-ACCAACGGAGGTCTTGTTGCAC-3’. For *odc*, F: 5’-CACATGTCAAGCCAGTTC-3’ and R: 5’-GCCTATACATTGATGCTG-3’.

## Acknowledgements

We thank Professors Shigeo Okabe and Akihiko Takashima for critical reading of the manuscript. This work was supported in part by grant from MEXT*-Supported Program for the Private University Research Branding Project 2016–2020 (*Ministry of Education, Culture, Sports, Science, and Technology).

## Abbreviations

AVE: anterior visceral endoderm
PFMH: prospective forebrain, midbrain, and hindbrain
VCC: vegetal cortical cytoplasm
WISH: whole-mount *in situ* hybridization

## Supplementary Information

Analysis of the relative mass of representative brain subdivisions in insectivores and dry-nosed primates shows that in a typical insectivore (median body weight 30g), the weight ratio of medulla: cerebellum: neocortex is 1.0 (0.063g): 0.8 (0.050g): 1.1 (0.069g), whereas in a typical dry-nosed primate (median body weight 3,000g), the ratio is 1.0 (1.29g): 3.9 (5.03g): 22.4 (28.9g). These values were calculated from published data (Barton, 2007). There appears to be a rostral bias in the magnification of the three subdivisions between insectivores and primates. The rostral bias was further confirmed at the cellular level. The ratio of number of neurons in medulla: cerebellum: cerebral neocortex is 1.0 (8.1×10^6^ cells): 8.2 (66.4×10^6^ cells): 1.7 (13.8×10^6^ cells) in a typical insectivore (median weight 30g), whereas the ratio is 1.0 (86.3×10^6^ cells): 33.6 (2,900×10^6^ cells): 13.5 (1,139×10^6^ cells) in a typical primate (median weight 3,000g). These values have been calculated from published data (Herculano-Houzel et al., 2007; Sarko et al., 2009). Thus, the magnitude of increase in both mass and number of neurons per representative brain subdivision in primates relative to insectivores is progressively greater rostrally (neocortex > cerebellum > medulla), with the ratio of magnitude of increase being 20.4: 4.9: 1.0 for mass and 7.9:: 1.0 for cell number.

---

**Fig. S1.**
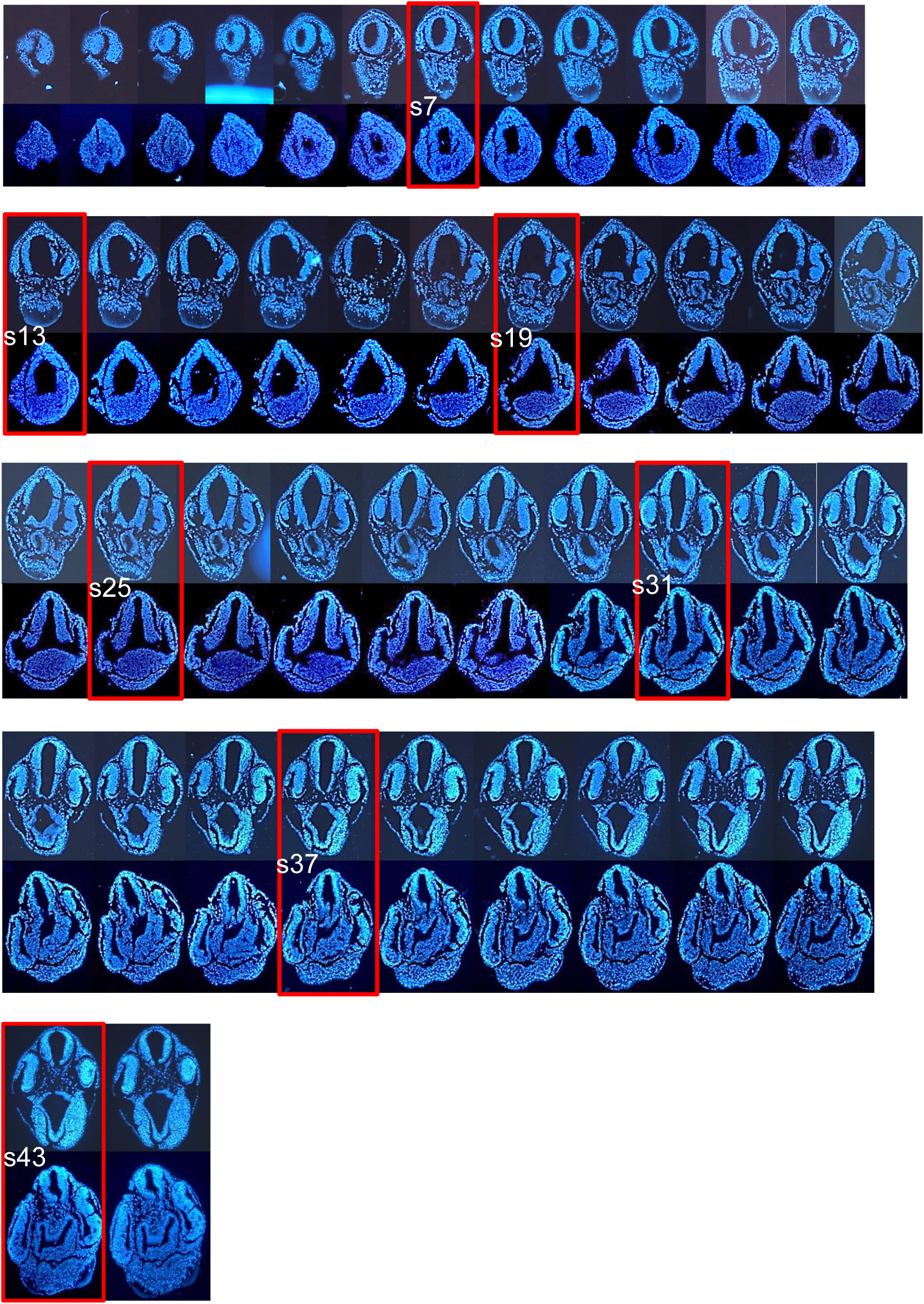
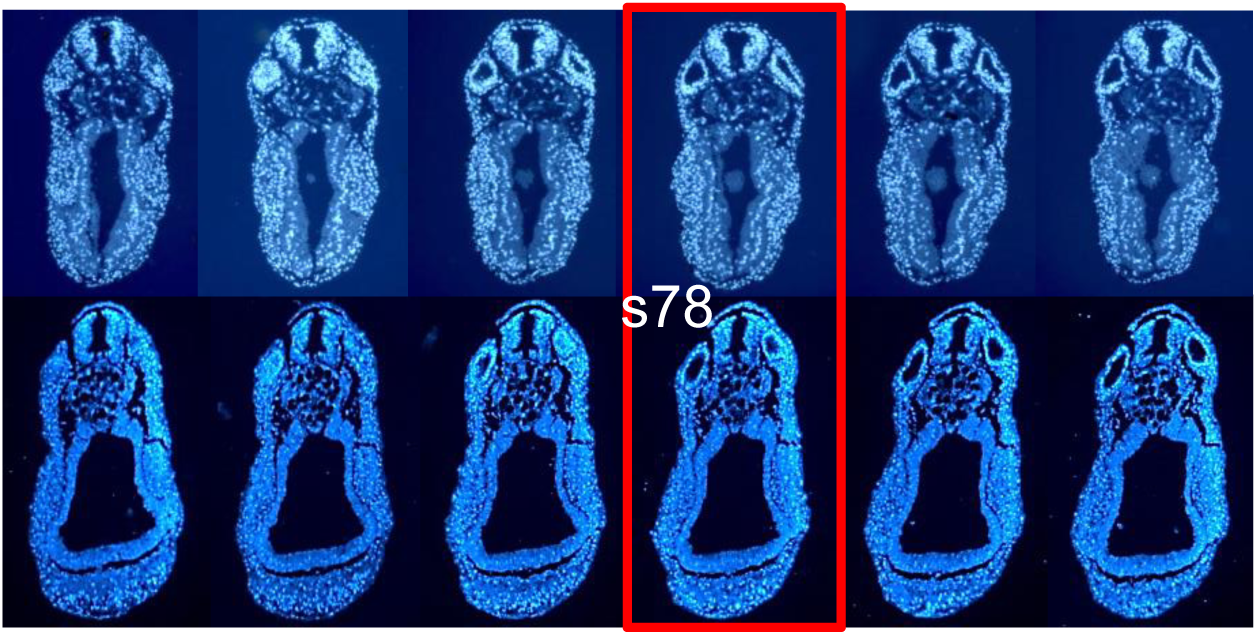
Supplementary data to Fig. 3. Consecutive series of sections are shown. The sections used in Fig. 3 are outlined in red. The upper sections are from a normal tailbud and the lower sections are from a Class I tailbud.

**Fig. S2.**
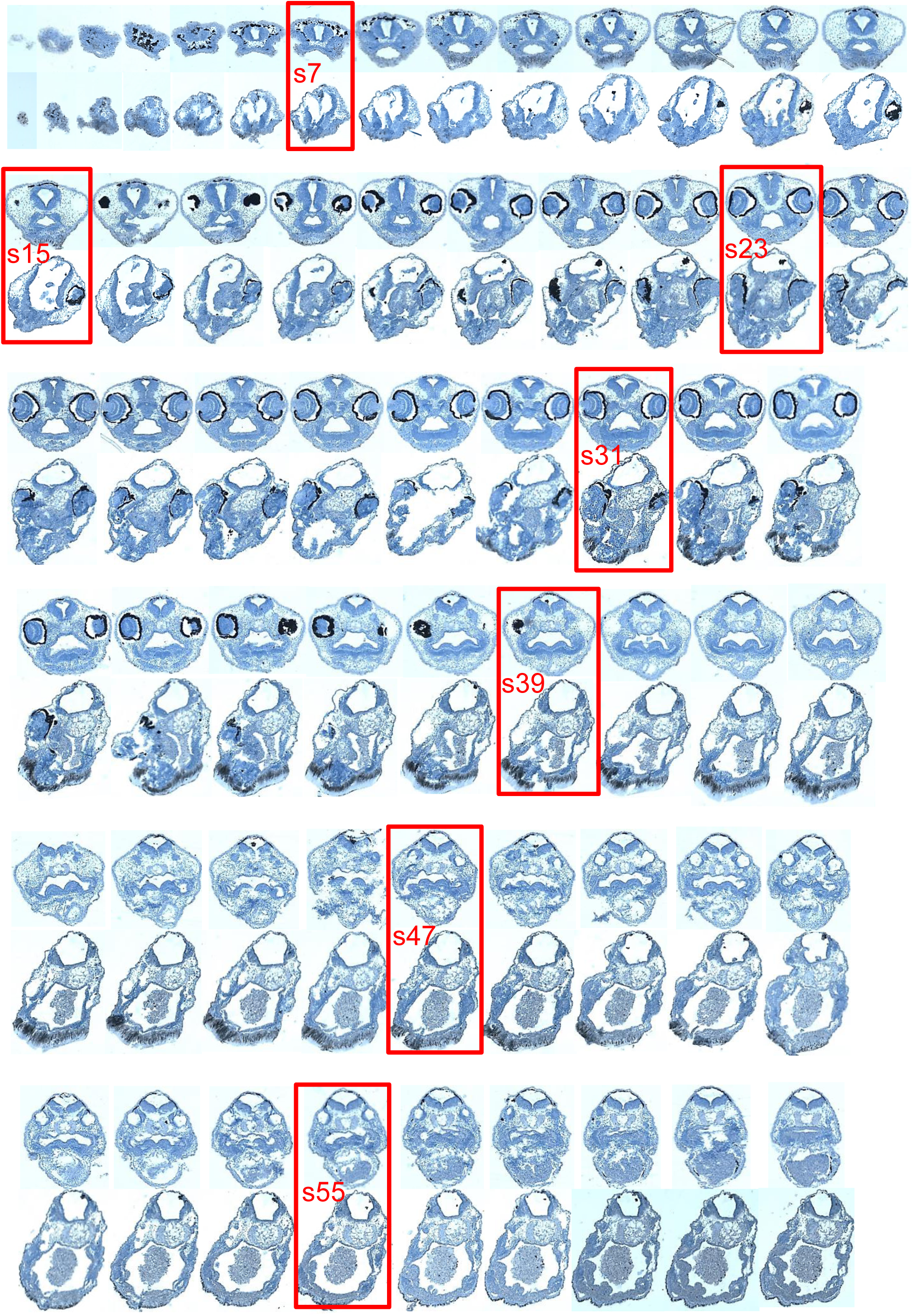
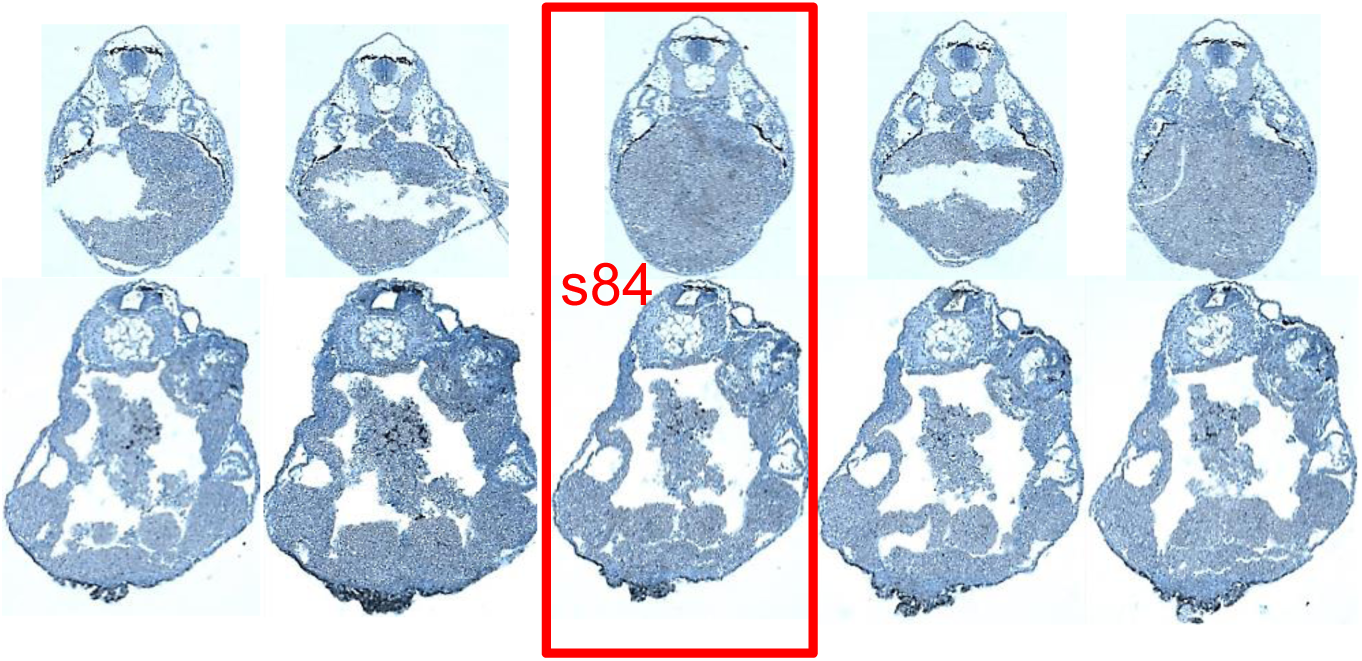
Supplementary data for Fig. 4. Consecutive series of sections are shown. The sections used in Fig. 4 are outlined in red. The upper sections are from a normal tadpole and the lower sections are from a Class I tadpole.

**Fig. S3.**
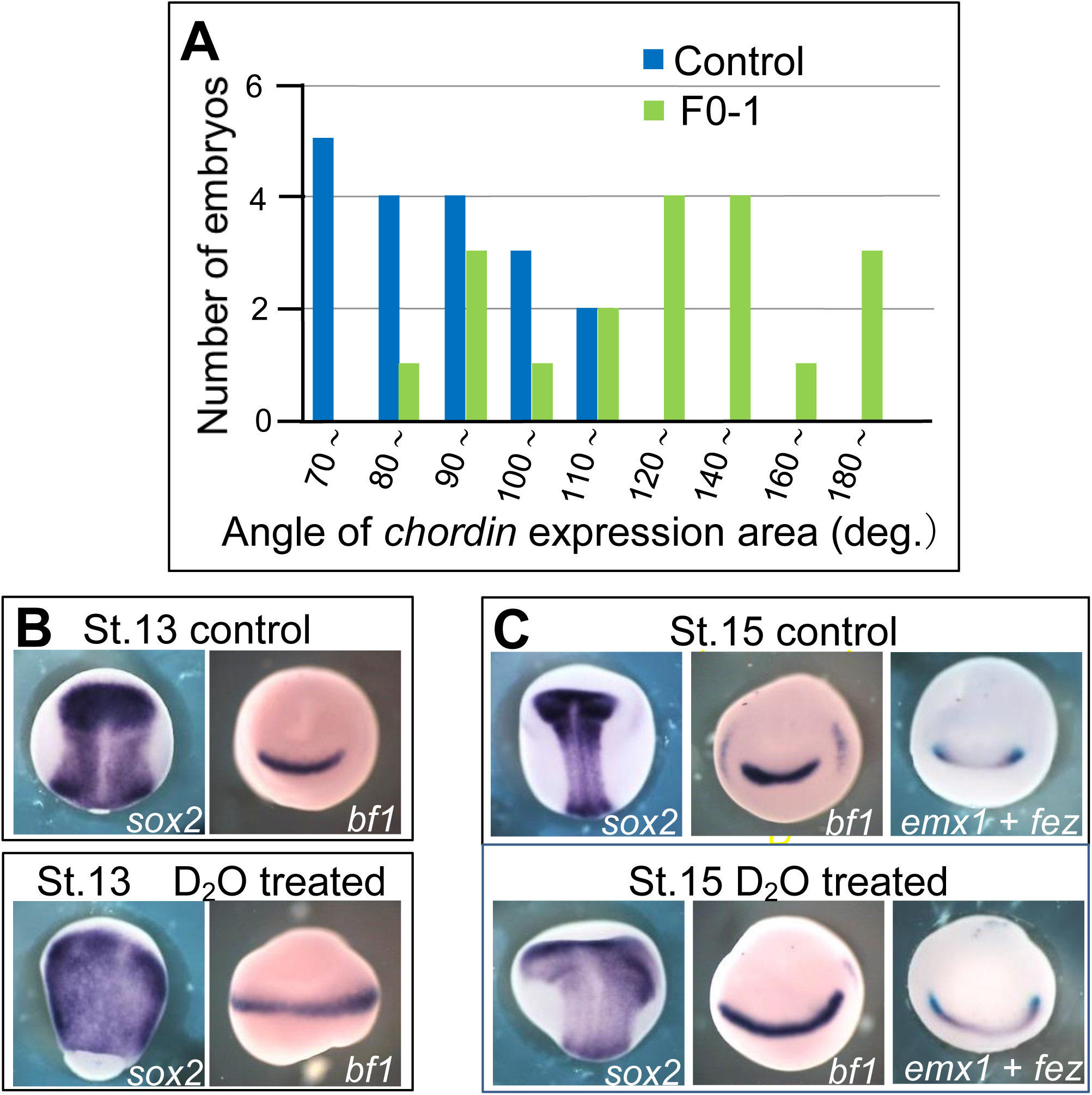
Expansion of the Speamann organiser and neural plate structures in D_2_O-treated embryos. Fertilised eggs from normal crosses were treated with D_2_O as reported (Scharf et al., 1989). (A) Expanded expression of *chordin* at stage 11 in D_2_O-treated embryos. Histograms were generated as described for Fig. 6A’. Mean ± SD values (degrees) for control and D_2_O-treated embryos are 90.1 ± 14.0 and 133.2 ± 333, respectively. The difference between the two groups is significant by t-test at p=3.6 × 10^−23^. (B) Expanded expression of *sox2* (left panels) and *bf1* (right panels) in D_2_O-treated embryos at stage 13. (C) Expanded expression of *sox2* (left panels), *bf1* (middle panels) and *emx1* + *fez* (right panels) in D_2_O-treated embryos at stage 15. *emx1* expression is light blue (BCIP) and *fez* expression is purple (BM purple). The double *in situ* procedures were essentially the same as described previously (Hongo and Okamoto, 2022).

**Fig. S4.**
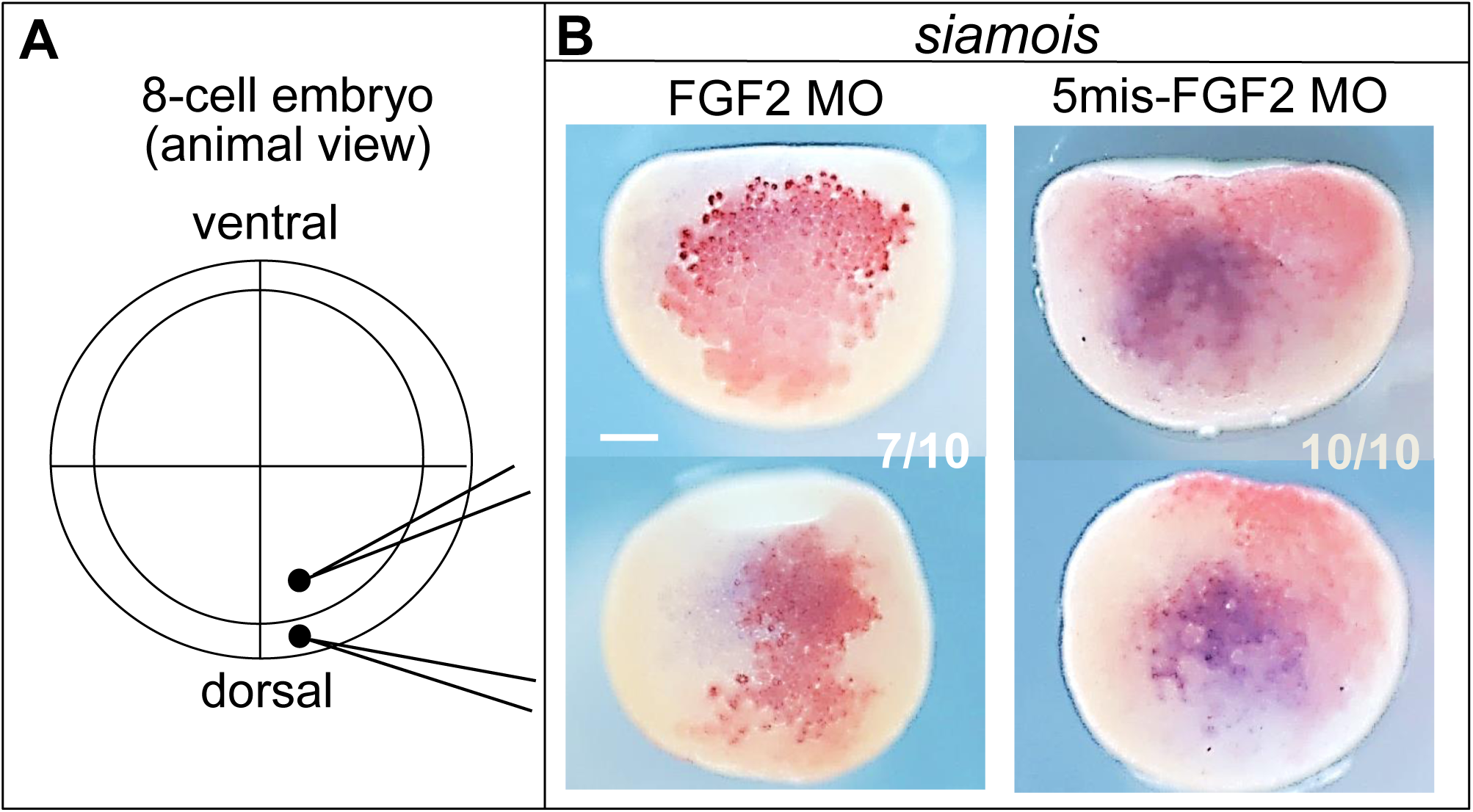
Suppression of *siamois* expression in late blastula by depletion of FGF2. (A) Injection protocol. Note that the injection site of MOs in both animal and vegetal blastomeres at the 8-cell stage was near the central equatorial region on the dorsal side of the embryo, so that the MOs would be distributed in the dorsal marginal zone of the embryo grown to the blastula stage. (B) FGF2 MO or 5mis-FGF2 MO was injected into a dorsal animal blastomere at a dose of 6.0 ng/blastomere and into a dorsal vegetal blastomere at a dose of 12.0 ng/blastomere, each co-injected with tracer *gfp* RNA (20 pg/blastomere). Whole-mount *in situ* hybridization was performed on blastula embryos at stage 9.5. The dorsal view is shown. The numbers on the panels indicate the number of embryos typically seen in the photographs relative to the total number of embryos analyzed. Scale bar = 0.20 mm. Microinjection and whole-mount double *in situ* hybridization were performed as previously described (Hongo and Okamoto, 2022).

